# Mutational screens highlight glycosylation as a modulator of CSF3R activity

**DOI:** 10.1101/2022.08.01.502088

**Authors:** Michael J. Hollander, Stacy A. Malaker, Idalia Perez, Julia E. Maxson, Jennifer R. Cochran, Carolyn R. Bertozzi

## Abstract

The colony-stimulating factor 3 receptor (CSF3R) controls the growth of neutrophils, the most abundant type of white blood cell. In healthy neutrophils, signaling is dependent on CSF3R binding to its ligand CSF3. A single amino acid mutation in CSF3R, T618I, instead allows for constitutive, ligand-independent cell growth and leads to a rare type of cancer called chronic neutrophilic leukemia (CNL). We investigated why this threonine to isoleucine substitution is the predominant mutation in CNL and how it leads to uncontrolled neutrophil growth. Using protein domain mapping, we demonstrated that the single CSF3R domain containing residue 618 is sufficient for ligand-independent activity. We then applied an unbiased mutational screening strategy focused on this domain and found that activating mutations are enriched at sites normally occupied by asparagine, threonine, and serine residues – the three amino acids which are commonly glycosylated. We confirmed glycosylation at multiple CSF3R residues by mass spectrometry, including the presence of GalNAc and Gal-GalNAc glycans at wild-type threonine 618. Using the same approach applied to other cell surface receptors, we identified an activating mutation, S489F, in the interleukin-31 receptor alpha chain (IL-31Rα). Combined, these results suggest a role for glycosylated hotspot residues in regulating receptor signaling, mutation of which can lead to ligand-independent, uncontrolled activity.

## Introduction

Cell surface receptors direct the first step of signaling pathways that govern many cellular activities^1^. Activation of receptor signaling commonly occurs when a ligand binds to the receptor. This binding event often changes the conformation or oligomerization of the receptor, thereby transmitting the signal across the cell membrane^2^. As a result, signaling proceeds in a controlled manner only in the presence of the ligand, allowing cells to maintain homeostasis. Examples exist of amino acid mutations which enable receptors to signal even in the absence of ligand, overriding this control mechanism^3^. This is especially notable for activating mutations in growth factor receptors which cause cells to proliferate uncontrollably, leading to several forms of cancer^4^.

Beyond amino acid substitutions, there is a need to consider the implications for altered posttranslational modifications in dysregulated growth factor receptor signaling. A predominant example is glycosylation, which typically modifies asparagine, serine, or threonine residues with complex glycans. N-linked glycosylation is named for sugar moieties attached to the nitrogen of asparagine in an Asn-X-Ser/Thr sequon, while O-linked glycosylation acts on the oxygen of serine or threonine^5^. Glycans are much larger than amino acid side chains, and can introduce negative charge through specific monosaccharides such as sialic acids, or through sulfate modifications^6^. As a result, when glycosylation sites are either introduced or removed by mutation, the effects can potentially be more consequential to overall protein structure than simple amino acid side chain substitutions.

A major challenge of studying protein glycosylation is difficulty in assigning the sites and structures of the attached glycans. Glycosylation is not genetically templated, so while it is possible for any threonine or serine to be glycosylated, the modification depends on the presence of enzymes, predominantly glycosyltransferases, that build glycans at specific amino acid residues^7^. Similarly, asparagine residues that are part of the defined glycosylation sequon may or may not be glycosylated depending on these enzymes and other factors^8^. When residues are glycosylated, there is additional macroheterogeneity – the fraction of sites that are occupied - and microheterogeneity – the range of glycan structures at individual sites^9^. Computational tools that predict glycosylation based on the sequence of a protein are still in development and require experimental validation^10^. We therefore may suspect that a threonine, serine, or asparagine residue is glycosylated but cannot have certainty without direct molecular characterization, typically by mass spectrometry-based glycoproteomic analysis^11^.

Colony stimulating factor 3 receptor (CSF3R, also G-CSFR) is an example of a cell surface growth factor receptor whose signaling activity was recently proposed to be regulated by glycosylation^12^. Wild-type CSF3R is a transmembrane protein that dimerizes in the presence of the ligand colony-stimulating factor 3 (CSF3) with a 2:2 ligand-receptor stoichiometry, leading to downstream JAK/STAT signaling and the development and proliferation of neutrophils, often in the context of infection where an innate immune response is beneficial^13,14^. CSF3R is comprised of an Ig-like domain at the N-terminus for ligand binding, five extracellular fibronectin-III (FnIII) domains, a transmembrane region, and an intracellular signaling domain (Figure 1A)^15^. Maxson et al. discovered that a cancer driving mutation, T618I in the 5^th^ FnIII (5FnIII) domain, is present in over 80% of cases of chronic neutrophilic leukemia (CNL), a myeloproliferative neoplasm characterized by high levels of mature neutrophils and band cells^16,17^. CNL is distinguished from other related malignancies by the presence of a CSF3R mutation and the absence of other genetic markers, such as the Philadelphia Chromosome. The average survival for patients with CNL is approximately 2 years after diagnosis^18^. Treatment options include hydroxyurea, bone marrow transplant, and the small molecule JAK inhibitor, ruxolitinib, which targets signaling downstream of CSF3R^19^.

**Figure 1.**
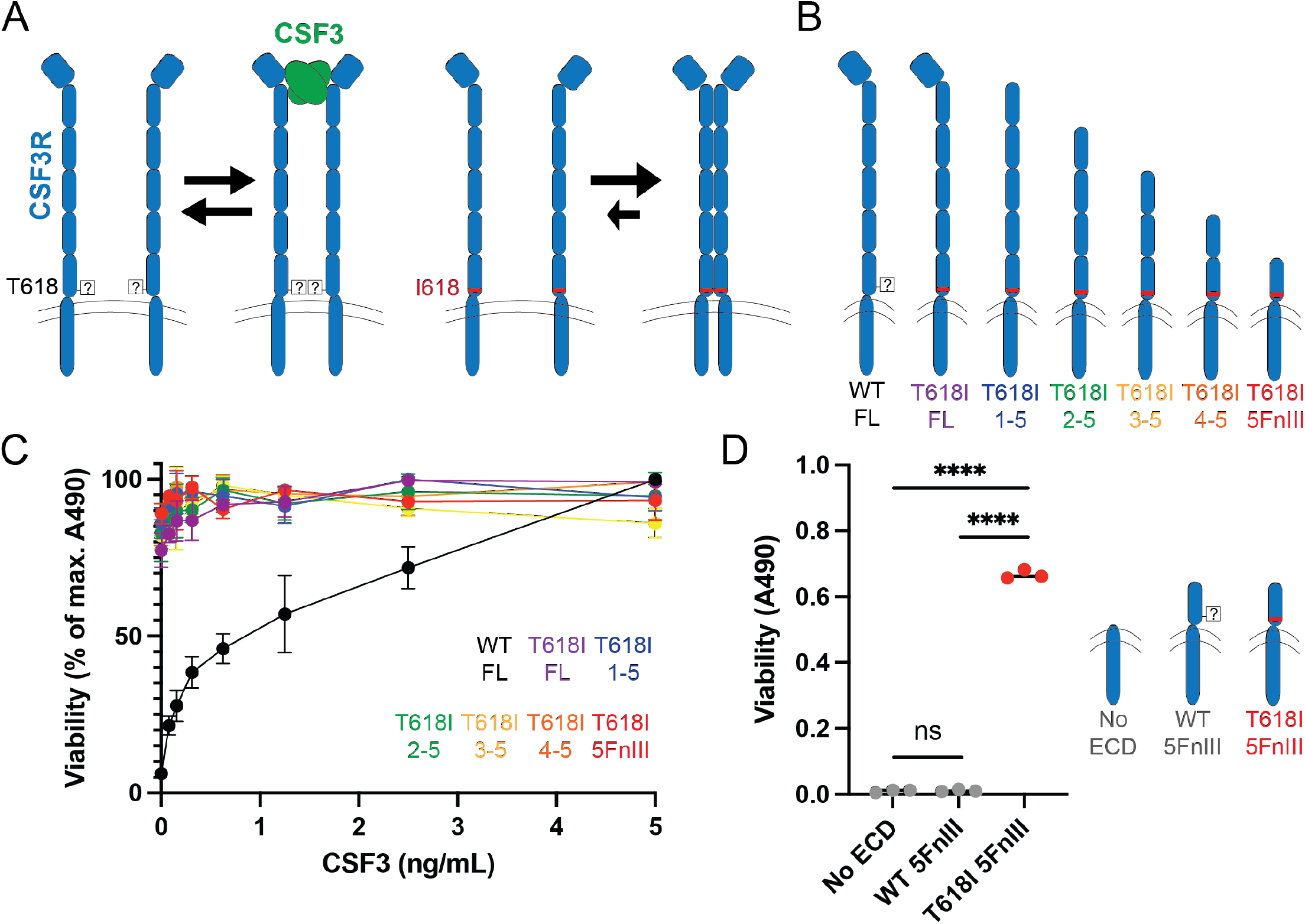
The membrane-proximal domain initiates ligand-independent signaling. A) Model for increased ligand-independent dimerization and signaling with the T618I mutation if it removes steric hindrance caused by a glycan at that site. B) Full-length and truncated constructs of CSF3R for determining which domain(s) are sufficient for activity. C) Dose-response curves for viability as measured by MTS assay. Values represent mean ± s.d. D) Comparison of mutant and WT 5^th^ FnIII domains and a control without the extracellular domain. Statistical significance was determined by unpaired t-test (****: *p* < 0.000001, ns: *p* = 0.64). Experiments were conducted at least two times with consistent results.

Maxson et al. demonstrated ligand-independent activity of CSF3R with the T618I mutation *in vitro* and showed that it recapitulates a CNL-like disease in a mouse model^20^. Furthermore, cells expressing the T618I mutant CSF3R proliferate even in the absence of CSF3 or other growth factor support. It was hypothesized that one key difference between wild-type threonine and the isoleucine substitution is that threonine can be modified by O-linked glycosylation, while isoleucine cannot; however, direct analysis of this putative glycosylation modification has not yet been reported. A more recently discovered leukemia-associated CSF3R mutation, N610H, does lead to a loss of N-linked glycosylation, as we recently determined using mass spectrometry methods^21^.

In this report, we performed an in-depth analysis of wild-type CSF3R to determine the effects of glycosylation on receptor signaling and cell growth. We identified the mutant membrane-proximal FnIII domain as the sole initiator of receptor signaling activity in the absence of the ligand CSF3. In addition, we performed a mutational screen for variants that induce CSF3R activation and found that a preponderance of activating mutations were ones that disrupted potential glycosylation sites. Finally, we expanded the screens to related receptors to discover whether these mechanisms are conserved beyond CSF3R. This study points to glycosylation of juxtamembrane domains of cell surface receptors as a regulatory modification that serve to curb aberrant activity.

## Results

### The 5^th^ FnIII domain of CSF3R is sufficient for cell viability

To probe how the T618I mutation controls CSF3R activity we sought to determine if the mutation has a local effect within the 5FnIII domain or global effects on the complete CSF3R conformation. A working model is that glycosylation could add steric hindrance at residue 618, whereas its removal by mutation could allow increased access for ligand-independent dimerization and signaling (Figure 1A). We expressed individual CSF3R domain fragments on the surface of mammalian cells to determine the minimal receptor regions required for activity. Mouse pro-B BaF3 cells are a model system for studying ligand-independent activity and have been applied to validate clinical mutations and screen for additional variants in mechanistic and biochemical studies^22,23^. Unlike most cultured cells which require only a base medium and serum for growth, BaF3 cells need either interleukin-3 (IL-3) supplementation to signal through endogenous IL-3 receptor (IL-3R), or an orthogonal signaling axis^24^. In this way, cells containing a signaling activating mutation have a growth advantage in media without IL-3.

We compared BaF3 activity for full-length wild-type and T618I CSF3R constructs to T618I variants with an increasing number of domains truncated from the N-terminus (Figure 1B). We observed the expected ligand-dependent response for full-length wild-type CSF3R, and ligand-independent viability for cells expressing fulllength T618I CSF3R. Interestingly, the 5FnIII T618I domain alone was sufficient for ligand-independent proliferation in an MTS-based viability assay (Figure 1C). The T618I mutation is critical, as the wild-type sequence of the 5FnIII construct does not confer activity. To determine whether intermolecular interactions stem from the 5FnIII or intracellular domain for T618I, we expressed a control containing only the transmembrane alpha helix and intracellular domain (residues 621-836) (Figure 1D). This control construct also did not yield viability, suggesting that interactions occur in the 5FnIII domain, and that the wild-type threonine is not permissive to these contacts. We therefore focused on the 5FnIII domain as the region of interest for mutational screens.

### Isoleucine substitution at 618 promotes hydrophobic contacts

It is unknown why the T618I mutation is most prevalent in CNL. Two variables are the specific residue location and the biochemical properties of the substituted amino acids. Previous work by Zhang et al. suggested that hydrophobic residues, like isoleucine, at position 618 promote ligand-independent signaling^25^. We therefore validated our high-throughput screen of CSF3R mutations by randomizing site 618 to allow any possible nucleotide combination at this position (termed an NNN codon). An advantage of this library design is that we can simultaneously test all 64 possible codons in one cell culture flask per condition. We cultured individual aliquots of the library under three conditions: 1) with mouse IL-3 to maintain the complete library as a starting point, 2) with human CSF3 to enrich for functional CSF3R variants that can signal in a ligand-dependent manner, and 3) with media without either ligand to enrich for activating mutations (Figure 2A). Following sequencing of the cultured libraries, we then calculated the fold-change enrichment by dividing the frequency in the ligand-independent group by the frequency in either the IL-3 or CSF3 group. As expected, hydrophobic and aromatic residues, most notably isoleucine, valine, phenylalanine, and tyrosine, are all enriched in the ligand-independent group (Figure 2B). Alanine and methionine, which are often grouped with nonpolar residues but are less hydrophobic, were not enriched. Interestingly, amino acids with similar properties clustered together when plotted for ligand-independent enrichment over IL-3 and CSF3. For example, charged amino acids (aspartic acid, glutamine acid, lysine, arginine, and histidine) all demonstrate similar fold changes. Threonine and serine, the two amino acids which can be O-glycosylated, are the most depleted.

**Figure 2.**
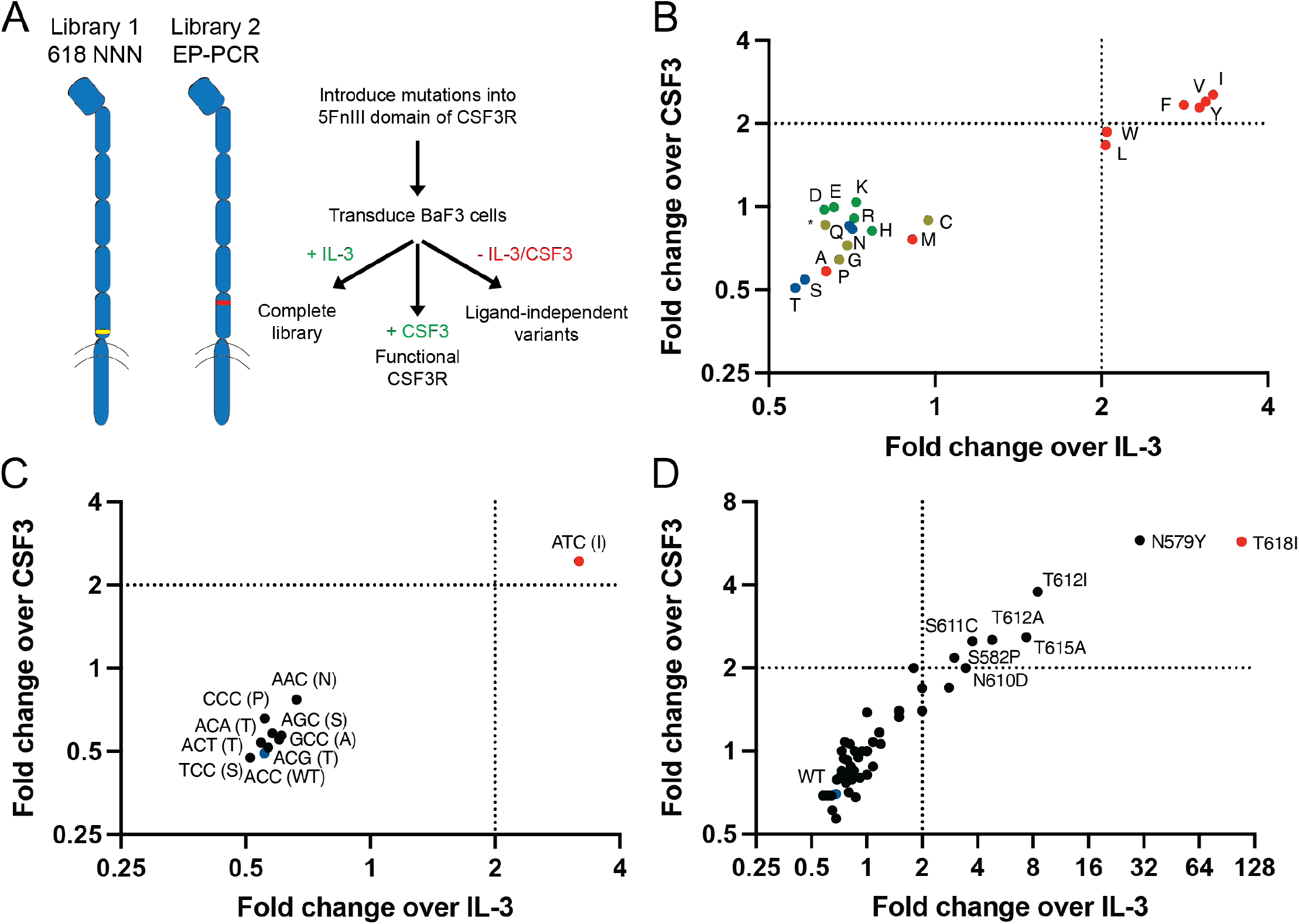
Enriched CSF3R mutations remove potential glycosylation sites. A) Workflow for CSF3R library design and screening. B) Substitutions at residue 618 from Library 1 (NNN codon), grouped by hydrophobic (red), charged (green), uncharged (blue), and other (yellow) amino acids. C) Possible codons at residue 618 with a single nucleotide mutation from the wild-type threonine (ACC, blue). D) The top 50 single nucleotide variants by frequency after screening Library 2 (error-prone PCR). Experiments were conducted at least two times with consistent results.

It is important to note that not all amino acid substitutions are possible with one nucleotide mutation at the codon for residue 618. Because we allowed for all possible codons, we could track codons individually for enrichment, including the nine possible single nucleotide variants from the wild-type ACC for T618. The clinically relevant T618I mutation results from a C to T transition (ACC to ATC). T618I may be prevalent in part because the C to T transition itself is more common in nature^26^, but also because ATC is the only codon out of 9 single-nucleotide mutations which encodes for a hydrophobic residue. The triplet ATC, which encodes isoleucine, is the only enriched codon in the ligand-independent group (Figure 2C).

### Glycosite mutations are enriched across the 5FnIII domain

We next asked why T618I, and to a lesser extent T615A and N610H, are more common than mutations at other positions in the 5FnIII domain of CSF3R. We created an unbiased library of variants by randomly introducing mutations in the 5FnIII domain by error-prone PCR, and again cultured cells with either IL-3, CSF3, or no ligand. While we expected to see enrichment of mutations at residue T618 due to its clinical relevance and its independent validation in oncogenesis, we strikingly observed that the most enriched sequences all removed a threonine, serine, or asparagine – the three amino acids with common glycosylation (Figure 2D). We did not observe such an effect for other residues which could shape protein conformation. Interestingly, one of the most enriched variants – N579Y – had not previously been found in patients at the time of the screen but has since been identified in a rare case of CNL^27^. We note that cells expressing wild-type CSF3R are found in the ligand-independent group, but are depleted compared to the IL-3 and CSF3 groups and arise due to random mutations in the genome resulting from the transduction process.

### Glycoproteomics confirms the modification of amino acid residues

Given the long-hypothesized model that T618I removes a glycosylation site along with the abundance of threonine, serine, and asparagine mutations in the error-prone library, we aimed to confirm glycosylation at T618 by mass spectrometry at these sites. Short peptides have high charge density and facilitate site-localization of glycans, but these peptides are challenging to create without standard protease (i.e. trypsin) cleavage sites near the residue of interest. We turned to the O-glycoprotease OpeRATOR which cuts N-terminally to threonine or serine residues if truncated O-linked glycosylation is present^28^. Using recombinant CSF3R protein expressed in human Expi293F cells, we first treated samples with sialidase to remove sialic acid and enhance the activity of OpeRATOR. Samples were then incubated with the OpeRATOR protease, as well as PNGase F to remove background glycosylation signal from N-linked glycans (Figure 3A). We chose to employ a higher-energy collision dissociation product dependent electron transfer dissociation (HCD-pd-ETD) instrument method. Here, HCD was used to first identify signature glycan oxonium ions, which then triggers ETD fragmentation on the same precursor mass. Since ETD does not rely on collision for fragmentation, the glycan remains intact and allows for site-specific localization of the glycan. Using this instrument method and manual validation, we confirmed the presence of the disaccharide Gal-GalNAc glycan at threonine 618 (Figure 3B). Other sugars, including the monosaccharide GalNAc at 618 and Gal-GalNAc at threonine 612, were identified across the CSF3R 5FnIII domain as well (Figure 3C). Depending on the presence of sialic acid, these glycans could be the (sialyl-)Tn or T antigens, respectively.

**Figure 3.**
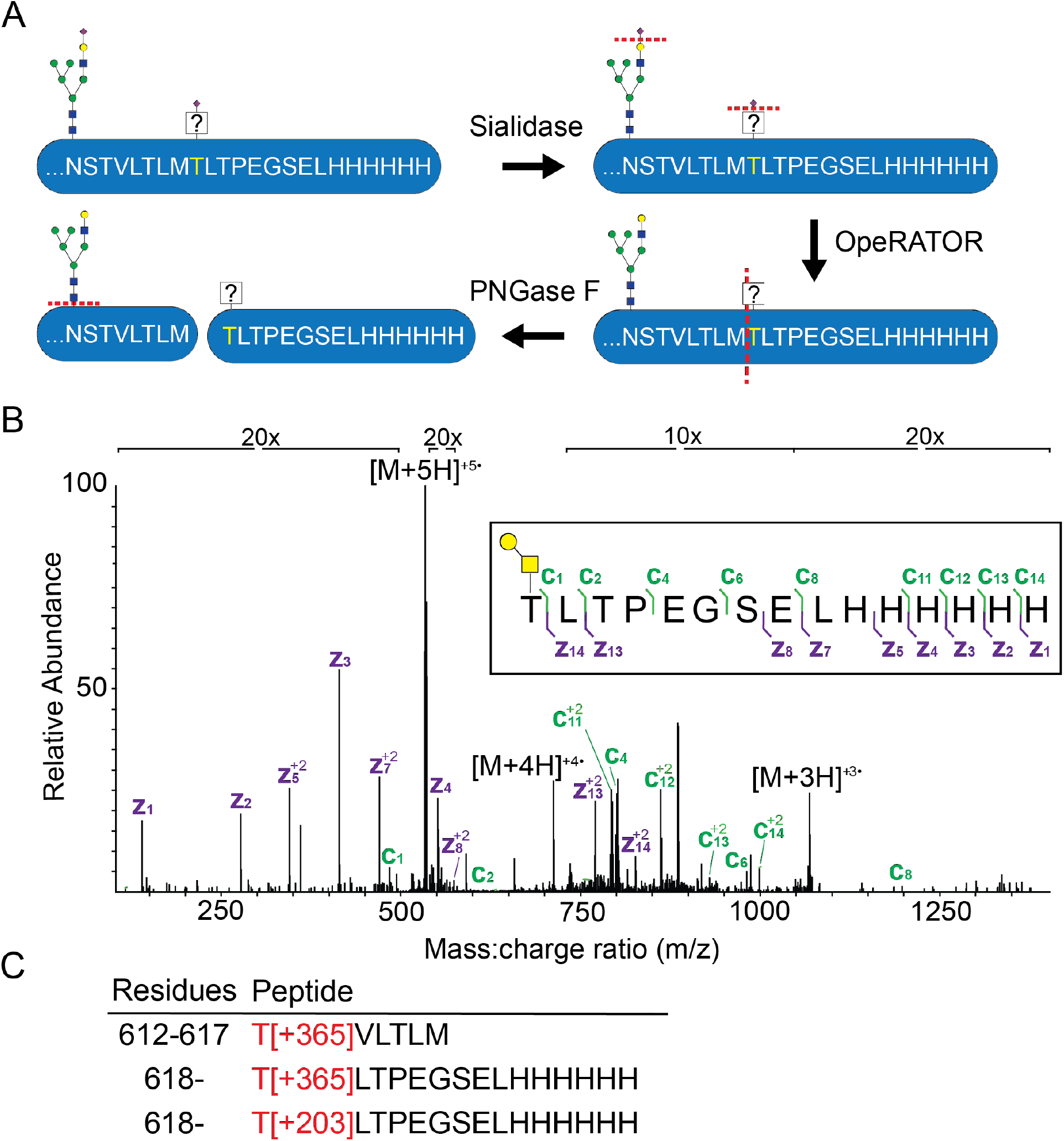
Wild-type threonine 618 is modified by glycosylation. A) Workflow for digesting a soluble form of the wild-type CSF3R 5FnIII domain with sialidase, the protease OpeRATOR, and PNGase F. B) Electron transfer dissociation (ETD) mass spectrum for the glycopeptide including the Gal-GalNAc (365 Da) glycan at T618. C) Observed glycopeptides throughout the CSF3R 5FnIII sequence, including the GalNAc (203 Da) glycan at T618 and Gal-GalNAc glycan at T612.

### Activating glycosite mutations are more common in CSF3R than related receptors

To determine how common such glycosite mutations are, we turned to proteins including the interleukin-6 (IL-6) family of receptors which have a similar structure to CSF3R. Of note, there is a high frequency of threonine and serine residues in the membrane-proximal FnIII domain of these receptors (Table 1). We selected proteins with an intracellular signaling domain and hydrophobic patches. We repeated the same library workflow to introduce random mutations into these domains and screen for ligand-independent BaF3 cell viability. For cells transduced with the single receptor libraries, only glycoprotein 130 (gp130) and leptin receptor (LEPR) cells expanded (Supplementary Table 1). The outgrowth of gp130 cells was due to a large deletion (Δ260-621) (Supplementary Figure 1). This was independently observed by rationally removing the stalk region^29^. LEPR cells expanded, but did not grow in a validation experiment (Supplementary Figure 2). Therefore, growth was likely due to a passenger mutation within the gene sequence and a driver mutation elsewhere in the genome. As such, these variants are not relevant in the context of this work.

**Table 1.**
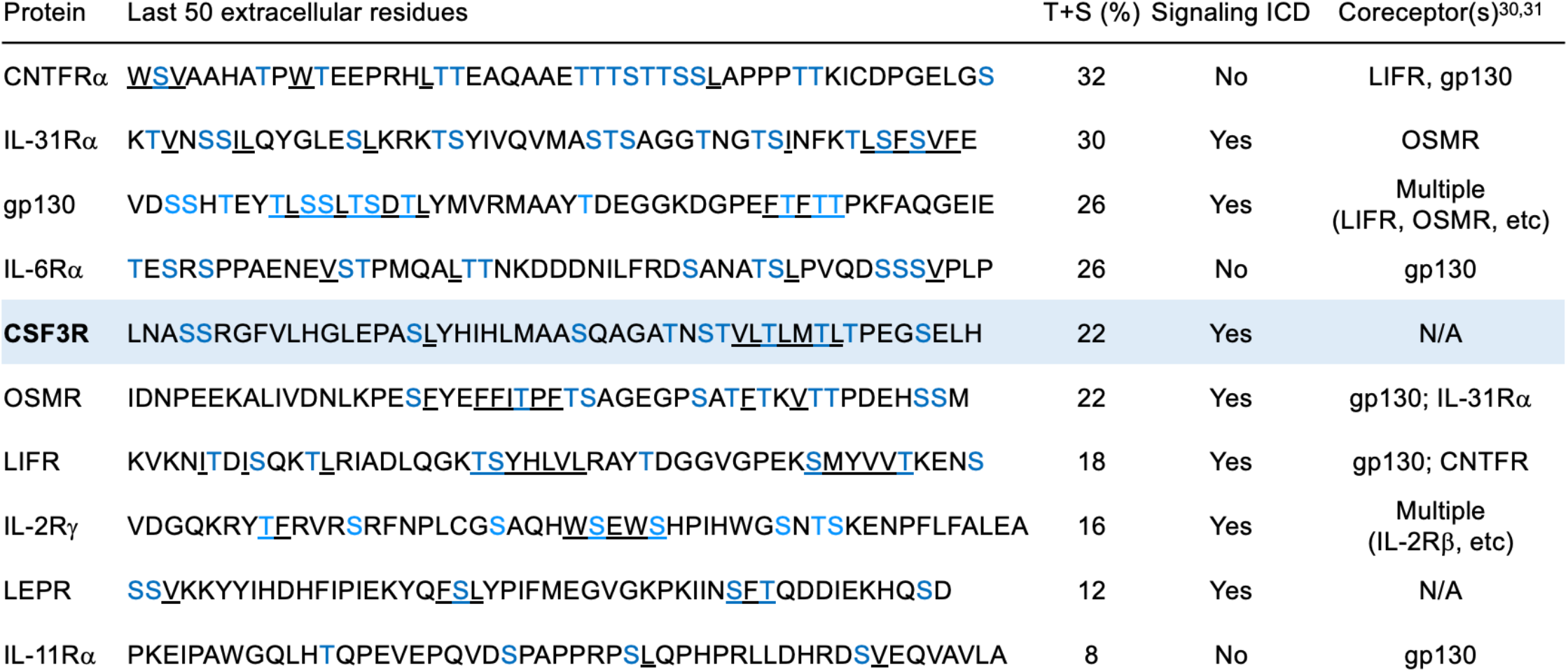
Cytokine receptors with high frequencies of threonine and serine residues in the membrane-proximal domain. Serine and threonine residues are highlighted in blue, and potential hydrophobic patches are underlined.

Many of these receptors typically form complexes with coreceptors for downstream signaling^30,31^, so we next repeated the screens by co-transducing the receptor library with its wild-type coreceptor: gp130/LIFR, LIFR/gp130, gp130/OSMR, OSMR/gp130, OSMR/IL-31Rα, and IL-31Rα/OSMR. Using this approach, we identified enriched clones including oncostatin-M receptor (OSMR) P734A variants with and without additional mutations (K705Q, F709L), as well as IL-31Rα variants (T427N, G433D, S489F, N509Y). We then validated these clones by transducing new BaF3 cells with the single mutations and growing them without ligand over time. Of the OSMR mutations, only P734A conferred ligand-independent growth in this outgrowth assay of BaF3 cells (Figure 4A). The other variants, K705Q and F709L, were therefore passenger mutations or unable to contribute to cell proliferation without another mutation. While P734 is predicted to cap the transmembrane alpha helix and mutations to that site are likely to change activity^32^, it is notable that many potential glycosites were not enriched in OSMR, and that the corresponding proline mutation in CSF3R, P621, was not enriched. Of the IL-31Rα mutations, S489F and N509Y were validated by outgrowth assay (Figure 4B). Although wild-type N509 is not part of a glycosylation sequon and cannot be modified by N-linked glycosylation, S489 may be modified by O-linked glycosylation. The S489F mutation (numbered S521F by Lin et al. due to a different isoform with 32 additional residues at the N-terminus, Supplementary Figure 3) is caused by a C to T transition like CSF3R T618I, and has been reported in a case of primary cutaneous amyloidosis^33^.

**Figure 4.**
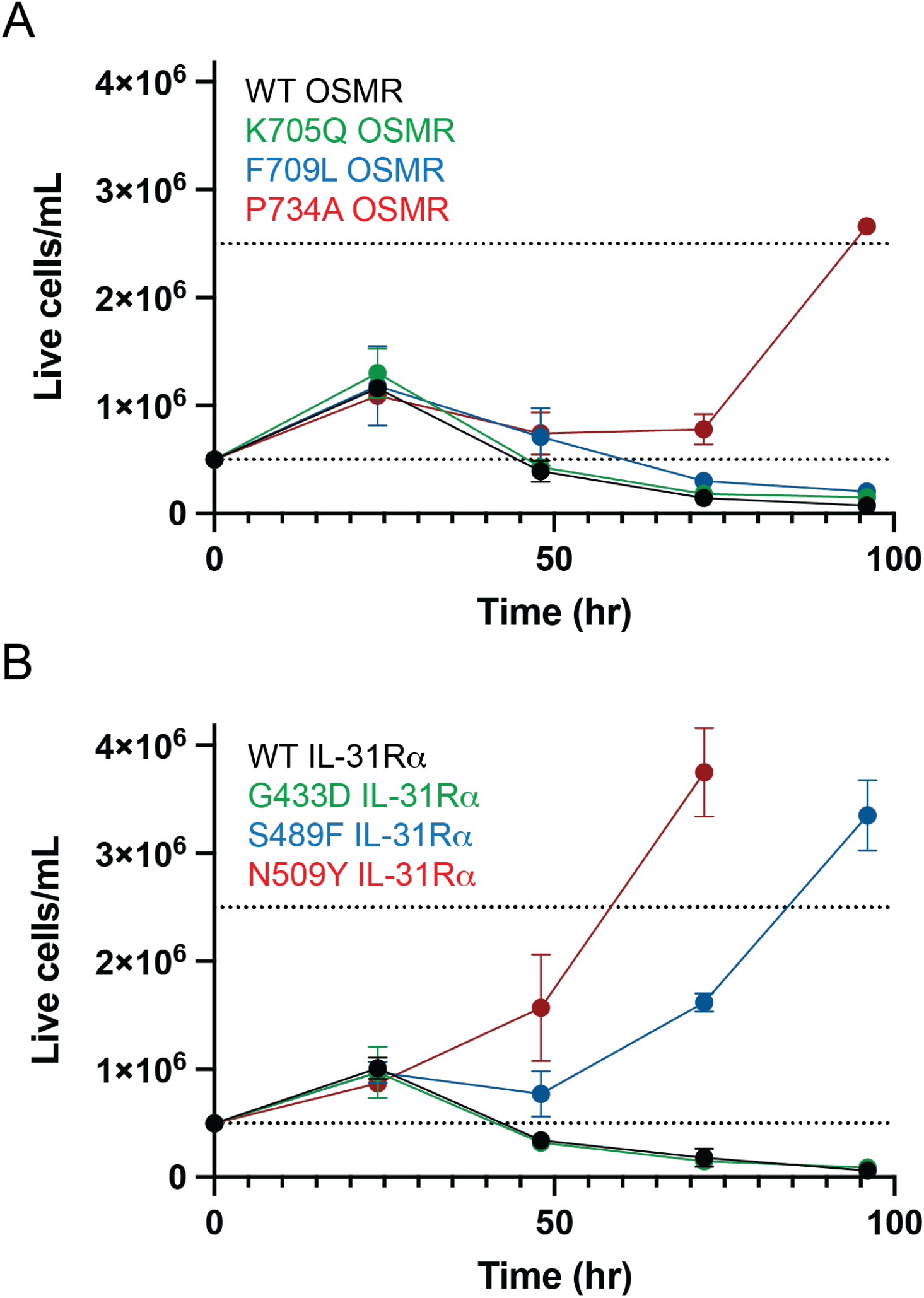
Activating mutations in related receptors. Outgrowth curves of BaF3 cells for OSMR variants transduced with wild-type IL-31Rα (A) and IL-31Rα variants transduced with wild-type OSMR (B). Values represent mean ± s.d.. Experiments were conducted at least two times with consistent results.

## Discussion

Studies of transmembrane receptors like CSF3R have often focused on ligand-binding domains, with only the three N-terminal domains represented in the published CSF3R crystal structure^34^. The membrane-proximal region of cell surface receptors is a growing focus of research in structural biology, investigating the role of intermediate protein domains and stalk regions between ligand-binding sites and the transmembrane region^35^. One example is the Tie2 receptor which, like CSF3R, was thought to dimerize in a ligand-dependent manner; however, an alternative ligand-independent mechanism was shown to be mediated by membrane-proximal FnIII domains^36^.

We focused on CSF3R based on clinically relevant mutations in CNL, and were particularly interested in the juxtamembrane domain given the results of the ligand-independent growth experiments (Figure 1) and the high frequency of serine and threonine residues which creates the possibility for clustered O-linked glycosylation, as encountered in so-called mucin-like domains^37^. Glycosylation has been shown to alter protein structure and function in multiple ways, including sterically occluding the underlying polypeptide as well as inducing specific polypeptide conformation^38,39^. Several groups have investigated the role of clustered O-glycans in either autolysis or cleavage by proteases, such as TNF-a by the metalloprotease ADAM17^40,41^.

Our findings instead highlight the consequences of removing glycosylation from proteins near the membrane. Using unbiased screens, we demonstrated that enriched activating mutations in the 5FnIII domain of CSF3R predominantly replace a potential glycosite with a hydrophobic residue, and that T618I is prevalent in part because of codon usage. Our findings indicate that membrane-proximal glycosylation serves as a “brake” to mediate receptor signaling. Rather than a binary “on/off” switch, the glycans allow for controlled signaling in the presence of ligand, and it is only when they are removed by mutation that aberrant signaling occurs. Of the seven receptors in this study, the phenomenon is most striking in CSF3R. One possible explanation is that CSF3R naturally forms a homodimer, rather than a heterodimer, in the presence of ligand. Consequently, any mutation could be found on both sides of the interface, which is especially important for the introduction of hydrophobic interactions. This is not the case for all homodimeric receptors, as we did not observe any mutations in LEPR that result in ligand-independent activity. For the receptors which form complexes with coreceptors, future studies could introduce mutations on both receptors at same time, though such combinations would be unlikely in patients.

It is also important to note that the exact positioning of these glycosites is key, as residues at the interface will likely impact signaling more than those on the non-interacting surfaces. Affinity is often thought to be driven by a combination of select “hotspot” residues and additional, weaker interactions from other amino acids^42^. Glycosylation may be similar, where sugar modification at key threonine, serine, or asparagine residues has an outsized effect on activity. When compared to IL-31Rα which also has an activating mutation at a potential glycosite, CSF3R may have a higher number of these glycosylation hotspots. As such, CSF3R may not necessarily be more prone to mutations in that domain, but when mutations arise, they could be more consequential. For this reason, full protein structures containing the membrane-proximal FnIII domains will provide additional insight into the mechanism of this activity.

In conclusion, this work confirms for the first time that there is O-linked glycosylation of CSF3R at wild-type T618. When mutated to isoleucine, we demonstrated that the 5FnIII domain containing residue 618 is sufficient for ligand-independent activity. Isoleucine is one of several hydrophobic residues which induce activation at this site, but the only one that can be reached with a single nucleotide mutation to the wild-type threonine codon. Beyond residue 618, we identified activating mutations at other sites within the 5FnIII domain, all of which remove potential glycosylation sites. Though similar mutations can be found in additional receptors like IL-31Rα, they are more common in CSF3R, likely due to the positioning of these hotspot glycosites. The clinical relevance of threonine and serine mutations extends beyond CSF3R and CNL to other instances where dysregulated signaling leads to disease. For example, the S489F mutation in IL-31Rα leads to primary cutaneous amyloidosis. With growing glycoproteomics and genomic variation datasets, there is an opportunity to further explore such mutations in the future.

## Experimental procedures

### DNA cloning for transmembrane receptor constructs

CSF3R constructs were cloned into an MSCV-IRES-GFP (MIG) plasmid digested with XhoI (New England Biolabs R0146S). For the constructs with truncated domains, two inserts were amplified by PCR using Phusion polymerase (New England Biolabs M0530S) according to the manufacturer’s protocols: the CSF3R signal peptide (residues 1-24), and the C-terminal region including FnIII domains 1-5 (residues 125-836), domains 2-5 (residues 233-836), domains 3-5 (residues 334-836), domains 4-5 (residues 431-836), domain 5 (residues 530-836), or the control without an extracellular domain (residues 621-836). These fragments were gel extracted (Thermo Scientific K0692) according to the manufacturer’s protocols, and added to Gibson assembly master mix (New England Biolabs E2611S). Stellar competent cells (Takara 636763) were transformed with the Gibson assembly products and spread on LB plates containing ampicillin. Individual colonies were miniprepped (Thermo Scientific K0503) and sequenced by Sanger sequencing (Elim Biopharm).

### Mammalian cell culture

HEK293T17 cells were cultured in DMEM (Fisher Scientific 11995073) supplemented with 10% fetal bovine serum (FBS, Thermo Fisher Scientific 26-140-079) and 1% penicillin-streptomycin (pen-strep, Fisher Scientific 15-140-122). BaF3 cells were cultured in RPMI 1640 (Fisher Scientific 11-875-119) supplemented with 10% FBS, 1% pen-strep, and 5 ng/mL recombinant murine interleukin-3 (IL-3, PeproTech 213-13-10ug). Expi293F cells (Thermo Fisher Scientific A14527) were cultured with shaking in Expi293 expression medium (Thermo Fisher Scientific A1435102). All cell lines tested negative for mycoplasma and were cultured at 37 °C in humidified incubators with 5% CO_2_ for HEK293T17 and BaF3 cells, and 8% CO_2_ for Expi293F cells.

### Transfection and transduction

CSF3R constructs were used to transfect HEK293T17 cells. First, 135 μL of Opti-MEM I reduced serum medium (Thermo Fisher Scientific 31985062) was mixed with 10 μL of FuGENE 6 transfection reagent (Promega E2693). After 5 minutes, 2 μg of receptor plasmid (1 μg each for co-receptors) and 0.83 μg of the EcoPac packaging vector were added, and after 15 minutes the complexes were dropped over a well of HEK293T17 cells in a 6-well plate. After 2 days, supernatant containing retrovirus for transductions was filtered with a 0.45 μm Steriflip (Fisher Scientific SE1M003M00).

For transduction, 1e6 BaF3 cells in 2 mL of complete media were mixed with 1 mL of media containing 10 ng/mL IL-3, 30 μL of 1M HEPES (Thermo Fisher Scientific 15-630-080), 2 μL of polybrene (Millipore Sigma TR-1003-G), and 1 mL of filtered supernatant containing retrovirus. Cells were centrifuged at 2500 rpm for 90 minutes at 30 °C with no brake and cultured overnight at 37 °C. On the next day, 2 mL were removed from the well, centrifuged at 1100 rpm for 5 minutes, resuspended with 1 mL of media containing 10 ng/mL IL-3, and added back to the cells along with an additional 30 μL of 1M HEPES, 2 μL of polybrene, and 1 mL of fresh supernatant containing retrovirus. Centrifugation was repeated with the same settings and cells were incubated overnight at 37 °C. On the following day, cells were centrifuged at 300xg for 5 minutes and resuspended in complete media without retrovirus. After expansion, transduced, GFP-positive cells were isolated by fluorescence-activated cell sorting (FACS) with an SH800 sorter (Sony), centrifuged at 300xg for 5 minutes, resuspended in complete media, and cultured at 37 °C.

### Ligand-independent viability assay

Sorted BaF3 cells cultured in complete media containing IL-3 were washed three times by centrifuging at 1200 rpm for 5 minutes and resuspending in 5 mL of complete media without IL-3. After the washes, cells were diluted to a density of 6e4/mL and 50 μL (3e3 cells) were added to 50 μL of media containing 2x of a serial dilution of human CSF3 (PeproTech 300-23) in a 96-well plate, for a total volume of 100 μL. After 3 days of incubation at 37 °C, 20 μL of CellTiter 96 AQueous One MTS reagent (Promega G3582) was added to cells. The absorbance at 490 nm was measured by a Synergy H4 plate reader (BioTek) according to the manufacturer’s protocols.

### Preparation of receptor libraries

For the randomized CSF3R library at residue 618, overlapping DNA oligos (IDT) were designed for the entire 5FnIII domain, with one containing the NNN codon at residue 618. The oligos were combined in a PCR assembly reaction (https://primerize.stanford.edu/protocol/#PCR). Ligations were used for cloning all libraries to maximize efficiency. Because SfiI, the restriction enzyme with a cut site closest to the 5FnIII domain, also cuts in the 3FnIII domain, the wild-type third and fourth FnIII domains were amplified by PCR. The 3-4FnIII fragment and randomized 5FnIII domain were digested with BsmBI-v2 (New England Biolabs R0739S), purified by PCR cleanup kit (Fisher Scientific FERK0702), combined with T4 ligase (Fisher Scientific 50-811-604), and then used as a template for PCR amplification of the complete insert. The full-length, wild-type CSF3R MIG plasmid and insert were digested with SfiI (New England Biolabs R0123S), ligated overnight with a 3:1 molar ratio, and finally purified by PCR cleanup kit. 25 μL of Endura electrocompetent bacteria (Lucigen 60242-1) were transformed with 150 ng of library DNA using a 0.1 cm Gene Pulser electroporation cuvette (Bio-Rad 1652083) and the following settings: 10 μF, 600 Ohms, 1800 V. The electroporated bacteria were then cultured in recovery medium (Lucigen 80026-1) for 1 hour, then 50 mL of LB with ampicillin overnight. On the following day, DNA was purified by midiprep kit (Fisher Scientific K210014).

For the error-prone CSF3R library, mutations were randomly introduced across the 5FnIII domain (residues 530-627) by adding 2.5 μL each of 20 μM 8-oxo-dGTP TriLink Biotechnologies N-2034) and 20 μM dPTP (TriLink Biotechnologies N-2037) to a 50 μL Taq polymerase (Fisher Scientific 50-811-694) reaction. The product was gel extracted and then used as a template for further PCR amplification. This amplicon was digested and ligated as described above for transforming bacteria.

For the gp130, LIFR, OSMR, IL-31Rα, LEPR, and IL-2Rγ libraries, eBlocks (IDT) were designed with restriction enzyme cut sites near the FnIII region of interest. Constructs for wild-type receptors were cloned into MIG by Gibson assembly as described above with the BstBI (Fisher Scientific 50-812-170) and NotI (Fisher Scientific 50-811-2040) restriction sites. After sequence verification, plasmids were digested with BamHI-HF (New England Biolabs R3136S)/MfeI-HF (New England Biolabs R3589S) (gp130), MfeI-HF/AgeI-HF (New England Biolabs R3552S) (LIFR), AgeI-HF/PsiI-v2 (Fisher Scientific NC1704764) (OSMR), BsiWI-HF (Fisher Scientific NC1240162)/BsaBI (Fisher Scientific 50-812-207) (IL-31Rα), HpaI (New England Biolabs R0105S)/AgeI-HF (LEPR), or BamHI-HF/PsiI-v2 (IL-2Rγ) and the backbones were purified by gel extraction. The juxtamembrane FnIII domain of each receptor (gp130: 518-619, LIFR: 724-833, OSMR: 625-740, IL-31Rα: 421-519, LEPR: 740-839, IL-2Rγ: 156-262) was mutagenized by error-prone PCR as described above, amplified, and digested with the respective enzyme pair. All library constructs were then ligated and transduced into BaF3 cells as previously described.

### Screens of receptor libraries in BaF3 cells

BaF3 libraries were washed three times as described above to remove IL-3 from the media. After the washes, cells were resuspended in 4 mL at a density of 0.5e6/mL in complete media containing 5 ng/mL IL-3, 5 ng/mL CSF3, or no ligand and cultured at 37 °C. On the following day, cells were split back to 4 mL at 0.5e6/mL. Cells were counted every day, and split to 4 mL at 0.25e6/mL after reaching a density greater than 1e6/mL. After cells were cultured for at least 1 week or until being passaged once, DNA was extracted from 2e6 cells using the QIAamp DNA Mini Kit (Qiagen 51304) according to the manufacturer’s protocols. The DNA encoding the FnIII domain of interest was amplified by PCR using primers with partial adapters for Amplicon-EZ (Azenta) analysis, purified by gel extraction, measured by a Qubit fluorometer (Thermo Fisher Scientific) and submitted to Azenta for next generation sequencing (NGS).

### Soluble protein expression and purification for mass spectrometry

The sequence for the wild-type CSF3R 5FnIII domain (residues 530-626) with the CSF3R signal peptide (residues 1-24) and a C-terminal 6xHis tag was cloned into a pAdd2 mammalian expression vector at the EcoRI and XhoI sites by Gibson assembly, as described above. Expi293F cells were transfected with the pAdd2 5FnIII construct using the ExpiFectamine 293 kit (Thermo Fisher Scientific A14524) according to the manufacturer’s protocols. 6 days after transfection, the supernatant was centrifuged at 300xg for 5 minutes, adjusted to a pH of 8 with NaOH, centrifuged again at 3700xg for 20 minutes, and filtered with a 0.22 μm bottle-top filter (EMD Millipore S2GPT02RE). His-tagged CSF3R protein was enriched by nickel-nitrilotriacetic acid (Ni-NTA) agarose (Qiagen 30210) and further purified by size-exclusion chromatography on a Superdex 75 10/300 GL column (Cytiva 17517401) with an AKTA Pure system (Cytiva).

### Mass spectrometry sample preparation and data analysis

After purification, 7 μg of wild-type 5FnIII protein was incubated overnight at 37 °C in a final volume of 30 μL with 7 U of both SialEXO sialidase (Genovis G1-SM1-020) and OpeRATOR (Genovis G2-OP1-20) enzymes diluted in 20 mM Tris pH 7.5. After 18 hours, the sample was diluted to 50 μL with ammonium bicarbonate buffer and incubated at 37 °C with 2 μL of PNGase F (New England Biolabs P0705S) diluted 1:100 in ammonium bicarbonate for an additional 7 hours. The sample was combined with 69 μL of ammonium bicarbonate along with 1 μL of 1% ProteaseMax surfactant (Promega V2072) for a total volume of 100 μL, and incubated at 25 °C overnight. After 18 hours, 0.3 μL of acetic acid was added to the sample before desalting with a C18 tip (Agilent A57203). The C18 tip was washed three times with 200 μL of methanol, then three times with 5% formic acid. The sample was applied to the tip for 30 seconds, then collected and reloaded for a total of 6 times. The tip was washed three times with 200 μL of 5% formic acid, then eluted three times with 100 μL of 80% acetonitrile and 5% formic acid. The sample was dried in a CentriVap Complete vacuum concentrator (Labconco) at 40 °C, resuspended in 5 μL of 0.1% formic acid, and frozen at −20 °C until analysis.

The frozen sample was analyzed as described by Shon et al.:

Samples were analyzed by online nanoflow liquid chromatography-tandem mass spectrometry using an Orbitrap Fusion Tribrid mass spectrometer (Thermo Fisher Scientific) coupled to a Dionex Ultimate 3000 HPLC (Thermo Fisher Scientific). A portion of the sample (4.5 μL of 5 μL; 40%) was loaded via autosampler isocratically onto a C18 nano precolumn using 0.1% formic acid in water (“solvent A”). For preconcentration and desalting, the column was washed with 2% acetonitrile and 0.1% formic acid in water (“loading pump solvent”). Subsequently, the C18 nano precolumn was switched in line with the C18 nano separation column (75-μm×250-mm EASYSpray containing 2μm C18 beads) for gradient elution. The column was held at 40 °C using a column heater in the EASY-Spray ionization source (Thermo Fisher Scientific). The samples were eluted at a constant flow rate of 0.3 μL/min using a 90-min gradient. The gradient profile was as follows (min:% solvent B, 2% formic acid in aceto-nitrile): 0:3, 3:3, 93:35, 103:42, 104:95, 109:95, 110:3, 140:3. The instrument method used an MS1 resolution of 60,000 full width at half maximum (FWHM) at 400 m/z, an automatic gain control (AGC) target of 3e5, and a mass range from 300 to 1,500 m/z. Dynamic exclusion was enabled with a repeat count of 3, repeat duration of 10 s, and exclusion duration of 10 s. Only charge states 2 to 6 were selected for fragmentation. MS2s were generated at top speed for 3 s. Higher-energy collisional dissociation (HCD) was performed on all selected precursor masses with the following parameters: isolation window of 2 m/z, 30% collision energy, orbitrap detection (resolution of 30,000), and an AGC target of 1e4 ions. Electron-transfer dissociation (ETD) was performed if 1) the precursor mass was between 300 and 1,000 m/z and 2) 3 of 9 HexNAc or NeuAc fingerprint ions (126.055,138.055, 144.07, 168.065, 186.076, 204.086, 274.092, and 292.103) were present at ±0.1 m/z and greater than 5% relative intensity. ETD parameters were as follows: calibrated charge-dependent ETD times, 2e5 reagent target, and precursor AGC target 1e4.
Raw files were searched using Byonic by Protein Metrics against directed databases containing the recombinant protein of interest. Search parameters included semi-specific cleavage specificity at the C-terminal site of R and K. Mass tolerance was set at 10 ppm for MS1s, 0.1 m/z for HCD MS2s, 0.35 m/z for ETD MS2s. Methionine oxidation (common 2), asparagine deamidation (common 2), and N-term acetylation (rare 1) were set as variable modifications with a total common max of 3, rare max of 1. O-glycans were also set as variable modifications (common 2), using the “O-glycan 6 most common” database. Cysteine carbaminomethylation was set as a fixed modification. Peptide hits were filtered using a 1% FDR. All peptides were manually validated and/or sequenced using Xcalibur software (Thermo Fisher Scientific). HCD was used to confirm that the peptides were glycosylated and ETD spectra were used for site-localization of glycosylation sites^43^.

### Outgrowth validation assay

BaF3 cells were transduced with receptor constructs for single mutations, as described above. After washing cells three times to remove IL-3, cells were resuspended in 4 mL of complete RPMI media without ligand at a density of 0.5e6/mL. Cells were counted every day until expanding five-fold to 2.5e6/mL. Extracted DNA was then re-analyzed by Sanger sequencing to confirm that no new mutations had been acquired.

## Data availability

The mass spectrometry proteomics data for Figure 3 have been deposited to the ProteomeXchange Consortium via the PRIDE partner repository^44^ with the dataset identifier PXD035664. All other raw data for Figures 1, 2, and 4 are included in the supporting information.

## Supporting information

This article contains supporting information.

## Acknowledgments

We thank Nicholas Riley, Gaelen Hess, Melissa Gray, and members of the Cochran and Bertozzi groups for advice and helpful discussions.

## Funding and additional information

Research reported in this publication was supported by the National Institutes of Health under Award Number F31CA243267 (to M.J.H.) and R01CA200423 (to C.R.B.), and by the Sarafan ChEM-H Predoctoral Training Program at the Chemistry/Biology Interface (to M.J.H.). S.A.M. was supported by a National Institutes of Health F32 Postdoctoral Fellowship (F32-GM126663-01) and is currently supported by the Yale Science Development Fund and a National Institutes of Health R35-GM147039. J.E.M. is supported by a Leukemia & Lymphoma Scholar Award, a Concern Foundation Conquer Cancer Now Award, an ACS Research Scholar Award and National Institutes of Health R01 HL157147. The content is solely the responsibility of the authors and does not necessarily represent the official views of the National Institutes of Health.

## References

1 Ventura, A. C. et al. (2014) Utilization of extracellular information before ligand-receptor binding reaches equilibrium expands and shifts the input dynamic range. Proc. Natl. Acad. Sci. U. S. A. 111, E3860–9

2 Gorby, C., Martinez-Fabregas, J., Wilmes, S. & Moraga, I. (2018) Mapping determinants of cytokine signaling via protein engineering. Front. Immunol. 9, 2143

3 Gómez, A., Wellbrock, C., Gutbrod, H., Dimitrijevic, N. & Schartl, M. (2001) Ligand-independent dimerization and activation of the oncogenic Xmrk receptor by two mutations in the extracellular domain. J. Biol. Chem. 276, 3333–3340

4 Valley, C. C. et al. (2015) Enhanced dimerization drives ligand-independent activity of mutant epidermal growth factor receptor in lung cancer. Mol. Biol. Cell 26, 4087–4099

5 Lee, H. S., Qi, Y. & Im, W. (2015) Effects of N-glycosylation on protein conformation and dynamics: Protein Data Bank analysis and molecular dynamics simulation study. Sci. Rep. 5, 8926

6 Wang, J.-R. et al. (2017) A method to identify trace sulfated IgG N-glycans as biomarkers for rheumatoid arthritis. Nat. Commun. 8, 631

7 Horvat, T., Zoldoš, V. & Lauc, G. (2011) Evolutional and clinical implications of the epigenetic regulation of protein glycosylation. Clin. Epigenetics 2, 425–432

8 Chang, D., Hackett, W. E., Zhong, L., Wan, X.-F. & Zaia, J. (2020) Measuring site-specific glycosylation similarity between influenza a virus variants with statistical certainty. Mol. Cell. Proteomics 19, 1533–1545

9 Ince, D., Lucas, T. M. & Malaker, S. A. (2022) Current strategies for characterization of mucin-domain glycoproteins. Curr. Opin. Chem. Biol. 69, 102174

10 Hansen, J. E. et al. (1998) NetOglyc: prediction of mucin type O-glycosylation sites based on sequence context and surface accessibility. Glycoconj. J. 15, 115–130

11 Riley, N. M., Malaker, S. A., Driessen, M. D. & Bertozzi, C. R. (2020) Optimal dissociation methods differ for N-and O-glycopeptides. J. Proteome Res. 19, 3286–3301

12 Maxson, J. E. et al. (2014) Ligand independence of the T618I mutation in the colony-stimulating factor 3 receptor (CSF3R) protein results from loss of O-linked glycosylation and increased receptor dimerization. J. Biol. Chem. 289, 5820–5827

13 Horan, T. et al. (1996) Dimerization of the extracellular domain of granuloycte-colony stimulating factor receptor by ligand binding: a monovalent ligand induces 2:2 complexes. Biochemistry 35, 4886–4896

14 Dwivedi, P. & Greis, K. D. (2017) Granulocyte colony-stimulating factor receptor signaling in severe congenital neutropenia, chronic neutrophilic leukemia, and related malignancies. Exp. Hematol. 46, 9–20

15 Zhang, H. et al. (2017) Unpaired extracellular cysteine mutations of CSF3R mediate gain or loss of function. Cancer Res. 77, 4258–4267

16 Maxson, J. E. et al. (2013) Oncogenic CSF3R mutations in chronic neutrophilic leukemia and atypical CML. N. Engl. J. Med. 368, 1781–1790

17 Pardanani, A. et al. (2013) CSF3R T618I is a highly prevalent and specific mutation in chronic neutrophilic leukemia. Leukemia 27, 1870–1873

18 Gotlib, J., Maxson, J. E., George, T. I. & Tyner, J. W. (2013) The new genetics of chronic neutrophilic leukemia and atypical CML: implications for diagnosis and treatment. Blood 122, 1707–1711

19 Dao, K.-H. T. et al. (2020) Efficacy of ruxolitinib in patients with chronic neutrophilic leukemia and atypical chronic myeloid leukemia. J. Clin. Oncol. 38, 1006–1018

20 Fleischman, A. G. et al. (2013) The CSF3R T618I mutation causes a lethal neutrophilic neoplasia in mice that is responsive to therapeutic JAK inhibition. Blood 122, 3628–3631

21 Spiciarich, D. R. et al. (2018) A novel germline variant in CSF3R reduces N-glycosylation and exerts potent oncogenic effects in leukemia. Cancer Res. 78, 6762–6770

22 Chakroborty, D. et al. (2019) An unbiased in vitro screen for activating epidermal growth factor receptor mutations. J. Biol. Chem. 294, 9377–9389

23 Jiang, J. et al. (2005) Epidermal growth factor-independent transformation of Ba/F3 cells with cancer-derived epidermal growth factor receptor mutants induces gefitinib-sensitive cell cycle progression. Cancer Res. 65, 8968–8974

24 Jeay, S., Sonenshein, G. E., Postel-Vinay, M. C. & Baixeras, E. (2000) Growth hormone prevents apoptosis through activation of nuclear factor-kappaB in interleukin-3-dependent Ba/F3 cell line. Mol. Endocrinol. 14, 650–661

25 Zhang, H. et al. (2018) Gain-of-function mutations in granulocyte colony–stimulating factor receptor (CSF3R) reveal distinct mechanisms of CSF3R activation. J. Biol. Chem. 293, 7387–7396

26 Lewis, C. A., Jr, Crayle, J., Zhou, S., Swanstrom, R. & Wolfenden, R. (2016) Cytosine deamination and the precipitous decline of spontaneous mutation during Earth’s history. Proc. Natl. Acad. Sci. U. S. A. 113, 8194–8199

27 Maniaci, B., Chung, J., Sanz-Altamira P., DeAngelo D. J. & Maxson J. E. (2021) A novel CSF3R activating mutation identified in a patient with chronic neutrophilic leukemia. Blood 138, 3582

28 Yang, S. et al. (2018) Deciphering protein O-glycosylation: Solid-phase chemoenzymatic cleavage and enrichment. Anal. Chem. 90, 8261–8269

29 Lamertz, L., Floss, D. M. & Scheller, J. (2018) Combined deletion of the fibronectin-type III domains and the stalk region results in ligand-independent, constitutive activation of the Interleukin 6 signal-transducing receptor gp130. Cytokine 110, 428–434

30 Lin, J.-X. & Leonard, W. J. (2018) The common cytokine receptor γ chain family of cytokines. Cold Spring Harb. Perspect. Biol. 10, a028449

31 West, N. R. (2019) Coordination of immune-stroma crosstalk by IL-6 family cytokines. Front. Immunol. 10, 1093

32 Nilsson, I. et al. (1998) Proline-induced disruption of a transmembrane alpha-helix in its natural environment. J. Mol. Biol. 284, 1165–1175

33 Lin, M.-W. et al. (2010) Novel IL31RA gene mutation and ancestral OSMR mutant allele in familial primary cutaneous amyloidosis. Eur. J. Hum. Genet. 18, 26–32

34 Tamada, T. et al. (2006) Homodimeric cross-over structure of the human granulocyte colony-stimulating factor (GCSF) receptor signaling complex. Proc. Natl. Acad. Sci. U. S. A. 103, 3135–3140

35 Kurth, I. et al. (2000) Importance of the membrane-proximal extracellular domains for activation of the signal transducer glycoprotein 130. J. Immunol. 164, 273–282

36 Moore, J. O., Lemmon, M. A. & Ferguson, K. M. (2017) Dimerization of Tie2 mediated by its membrane-proximal FNIII domains. Proc. Natl. Acad. Sci. U. S. A. 114, 4382–4387

37 Malaker, S. A. et al. (2022) Revealing the human mucinome. Nat. Commun. 13, 3542

38 Walls, A. C. et al. (2016) Glycan shield and epitope masking of a coronavirus spike protein observed by cryo-electron microscopy. Nat. Struct. Mol. Biol. 23, 899–905

39 Shurer, C. R. et al. (2019) Physical principles of membrane shape regulation by the glycocalyx. Cell 177, 1757–1770.e21

40 Remacle, A. G. et al. (2006) O-glycosylation regulates autolysis of cellular membrane type-1 matrix metalloproteinase (MT1-MMP). J. Biol. Chem. 281, 16897–16905

41 Goth, C. K. et al. (2015) A systematic study of modulation of ADAM-mediated ectodomain shedding by sitespecific O-glycosylation. Proc. Natl. Acad. Sci. U. S. A. 112, 14623–14628

42 Thanos, C. D., DeLano, W. L. & Wells, J. A. (2006) Hot-spot mimicry of a cytokine receptor by a small molecule. Proc. Natl. Acad. Sci. U. S. A. 103, 15422–15427

43 Shon, D. J. et al. (2020) An enzymatic toolkit for selective proteolysis, detection, and visualization of mucin-domain glycoproteins. Proc. Natl. Acad. Sci. U. S. A. 117, 21299–21307

44 Perez-Riverol, Y. et al. (2019) The PRIDE database and related tools and resources in 2019: improving support for quantification data. Nucleic Acids Res. 47, D442–D450

