## Supporting Information (Table S1, Figures S1-3) for "Mutational screens highlight glycosylation as a modulator of CSF3R activity"

| Receptor library | Enriched mutations | Validated by outgrowth |
| --- | --- | --- |
| gp130 | Δ260-621 | Yes |
| LIFR | None | N/A |
| OSMR | None | N/A |
| IL-31Rα | None | N/A |
| LEPR | D839EfsX6 | No |
| IL-2Rγ | None | N/A |

| Co-receptor library | Enriched mutations | Validated by outgrowth |
| --- | --- | --- |
| gp130 + WT LIFR | None | N/A |
| LIFR + WT gp130 | None | N/A |
| gp130 + WT OSMR | None | N/A |
| OSMR + WT gp130 | None | N/A |
| OSMR + WT IL-31Rα | OSMR K705Q | No |
| | OSMR F709L | No |
| | OSMR P734A | Yes |
| IL-31Rα + WT OSMR | IL-31Rα G433D | No |
| | IL-31Rα S489F | Yes |
| | IL-31Rα N509Y | Yes |

**Supplementary Table 1** – Summary of mutations from (co)receptor libraries.

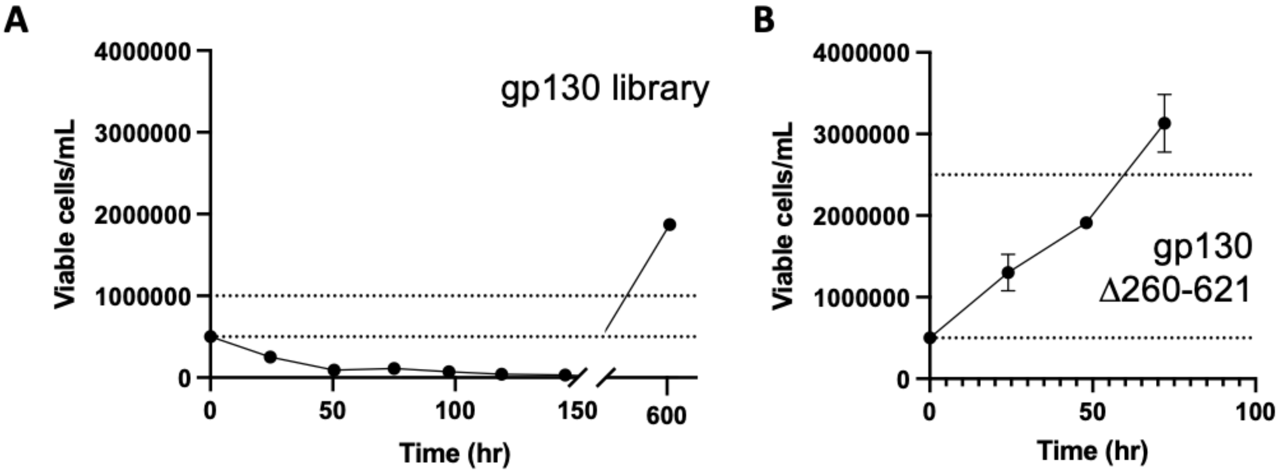


**Supplementary Figure 1** – Screen of gp130 library (A) and validation on enriched gp130 Δ260-621 clone by outgrowth assay.

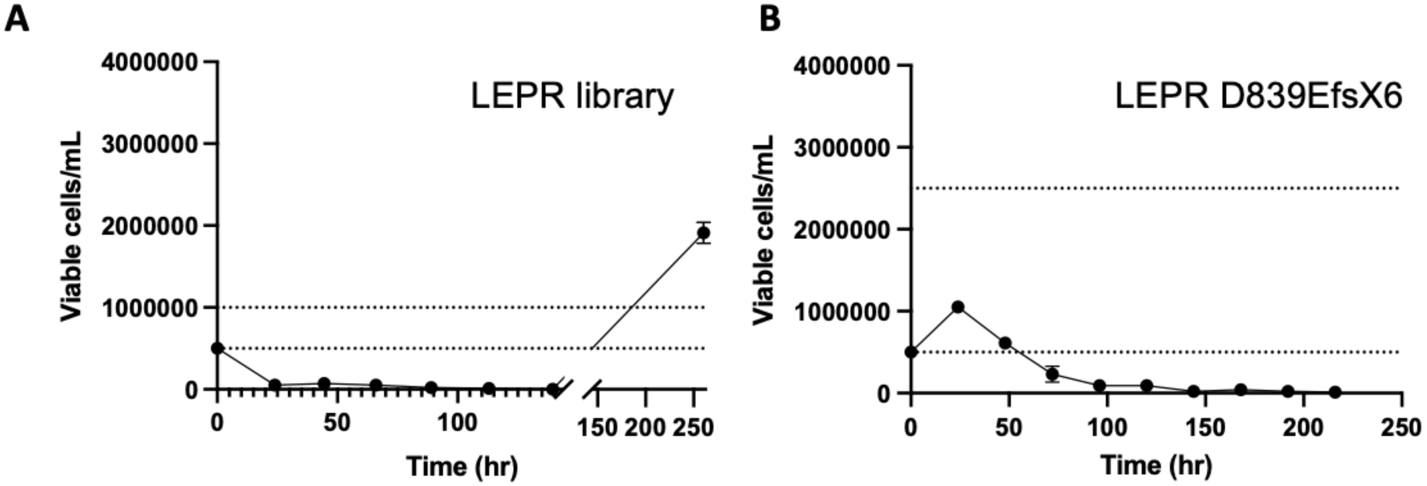


**Supplementary Figure 2** – Screen of LEPR library and outgrowth test for enriched frameshift mutation, LEPR D839EfsX6.

| | | | | | | |
|---|---|---|---|---|---|---|
| CGA | AAG | ACC | TCT | TAC | ATT | GTT |
| R486 | K487 | T488 | S489 | Y490 | I491 | V492 |

| | | | | | | |
|---|---|---|---|---|---|---|
| CGA | AAG | ACC | TTT | TAC | ATT | GTT |
| R486 | K487 | T488 | F489 | Y490 | I491 | V492 |

**Supplementary Figure 3** – Alignment of canonical IL-31Rα sequence to the region featured in Figure 3 by Lin et al.[33] The NM_139017 isoform includes 32 additional residues (MCIRQLKFFTTACVCECPQNILSPQPSCV… NLG) at the N-terminus. Consequently, S489 corresponds to S521.
